# A xylem-specific transcriptional atlas identifies ZHD transcription factors as central transmitters of protoxylem morphology

**DOI:** 10.64898/2026.11.13.766425

**Authors:** Akshay Gokulendran Nair, Claudia von der Mark, Jasper Staut, Thomas Depuydt, Thomas Eekhout, Carolin Grones, Maite Saura-Sanchez, Max Minne, Elisa Drijvers, Jonah Nolf, Jos Wendrich, Annika Saß, Ulrike Lehmann, René Schneider, Klaas Vandepoele, Bert De Rybel

## Abstract

Xylem cell development is characterized by deposition of a rigid secondary cell wall followed by programmed cell death leading to reinforced hollow tubes enabling water transport and providing physical strength. The molecular network controlling this differentiation pathway is well established, but the early events remain poorly understood. Here, we generated and extensively validated a high resolution, xylem-specific single cell RNA-sequencing atlas of the developing xylem. Subsequent gene regulatory network inference of this dataset identified the transcription factors *ZINC FINGER HOMEODOMAIN 3* (*ZHD3*) and *ZHD4* as central regulators of protoxylem development. We show that ZHD3/4 are required for normal protoxylem development and sufficient to trigger protoxylem-like SCW depositions in other xylem lineages by transcriptionally activating the xylem differentiation pathway. This identifies ZHD3/4 as central transmitters of PX morphology and questions whether SCW morphology serves as a valid determinant of cellular identity.

## Introduction

Vascular tissues feature a great diversity in function, morphology and development. Besides their role as conducting tissues, vascular cells serve as meristematic pools, storage sites or provide structural support to the plant body. Xylem cells transport water and minerals throughout the plant. In the primary root meristem of *Arabidopsis thaliana*, five xylem cells align to form the xylem axis at the center of the vascular cylinder (Fig. 1A). These arise from three xylem initials, which are located right above the quiescent center (ǪC). Two flanking initials will undergo one round of periclinal division to generate the protoxylem (PX) and outer metaxylem (MX), the third initial becomes inner MX [1] (Fig. 1A).

**Figure 1.**
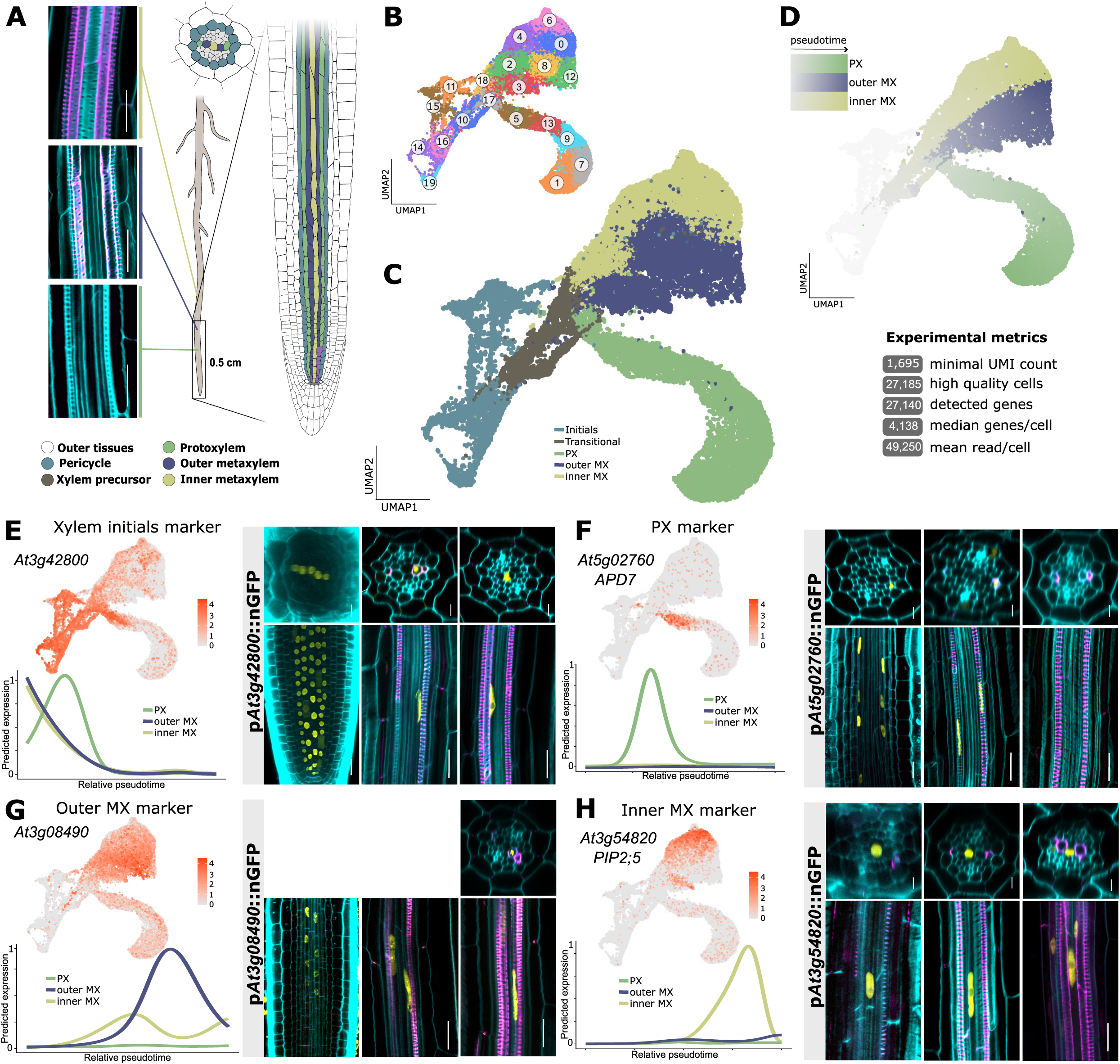
Xylem subpopulations display distinct transcriptomic landscapes. **A.** Schematic overview of xylem development in the Arabidopsis primary root meristem showing the morphological features and timing of differentiation associated with each xylem cell file. **B.** UMAP representation of the scRNA-seq dataset showing the initial clustering. **C.** Final annotated and validated UMAP including experiment metrics. **D.** Pseudotime analysis of the scRNA-seq data for the three xylem lineages. **E-H**. Example marker gene for each of the different xylem lineages (initials, PX, OMX and IMX) including the feature plot (top left), expression over pseudotime (bottom left) and confocal validation (right) showing both cross sections and longitudinal sections of the root apical meristem. Three positions in the root meristem are shown covering the longitudinal developmental zones: first appearance of the fluorescent signal, lignin deposition in PX and in outer MX. Cell walls are counterstained using Calcofluo white (cyan), lignin is stained using basic fuchsin (magenta), and gene expression (p*GENE*::nGFP) in yellow. Scale bars are 20 μm.

Mature xylem vessels, also named tracheary elements, are characterized by rigid secondary cell walls (SCW) which support the stability of the plant and withstand the hydraulic pressure within the vessel. Besides their localization along the xylem axis and developmental origin, PX and MX also show different patterns of SCW deposition. PX is characterized by a helical cellulose and lignin deposition while fortification in MX forms a pitted pattern [2,3]. Another aspect of xylem differentiation is developmental programmed cell death (dPCD). During this process all cellular content is degraded, leaving behind a hollow, rigid tube optimized for long-distance water transport. Both SCW deposition and execution of dPCD are enforced through a regulatory network with *VNDC* and *VND7* as central regulators [4–7]. *VNDs* activate, directly or indirectly through members of the MYB transcription factor family (most prominently *MYB4C* and *MYB83* [8,9] SCW cellulose synthases (*CESA4*, *CESA7*, and *CESA8*) or genes facilitating lignification like laccases and peroxidases [10–12]. As CESA complexes migrate along the microtubular network, the aforementioned characteristic patterns of helical PX and pitted MX cell walls are a consequence of microtubule rearrangements [13,14].

Despite extensive knowledge about the downstream processes of xylem differentiation following *VND* activation [4,5,15], little is known about early developmental regulators acting before this well characterized differentiation pathway. This is, in part, due to the fact that developing xylem cells are scarce, hidden in the most inner part of the root and characterized by rapid changes in physiology, progressing from meristematic initials to specified and fully mature xylem within a short time frame. These developmental trajectories are dictated by changes in the transcriptomic landscape of the maturing cell as it progresses through the root meristem. This high spatio-temporal complexity in the development of xylem cells thus demands a tool with equally high resolution, allowing the identification of early regulators of xylem development.

## Results

### Xylem subpopulations display distinct transcriptomic landscapes

Given that xylem cells are lowly abundant in Arabidopsis primary root meristems and difficult to isolate using cell digestion protocols, xylem cells make up less than 5% in published scRNA-seq atlases [16,17]. This complicates subsequent downstream computational analyses and hinders gene discovery pipelines. To eliminate this issue of underrepresentation, we sampled available xylem-specific reporter lines (p*ATHB8*::YFP, p*ACL5*::GFP, and p*TMO5*::tdTomato) (Fig. 1A, S1A) and enriched for xylem cells prior to scRNA-seq library preparation using Fluorescence-Activated Cell Sorting (FACS) (Fig. S1A). This multi-reporter strategy enabled the comprehensive capture of diverse xylem cell subtypes across multiple developmental stages. Low quality cells were removed followed by cell-type clustering and annotation leveraging expression patterns of known cell-type marker genes (Table S1). As some of the used reporters also marked non xylem cells (e.g. ATHB8 in columella cells, Fig. S1A) these clusters were removed from the data set based on marker gene analysis. This resulted in a final dataset containing 27,185 high quality xylem cells, expressing 27,140 genes in total (Fig. 1D, E). Clusters were computationally annotated by label transfer (see materials and methods for details) and cell type specific marker genes were manually checked to confirm the annotation and developmental lineage (Fig. S2). To validate these predicted cluster annotations, we next generated 23 reporter lines selecting differentially expressed genes (DEGs) from each cluster (Fig. 1E-H, Fig, S3). In all cases, expression of the reporter line followed the predictions from the scRNA-seq dataset, confirming the validity of the cluster annotations and the predictive power of the dataset (Fig. 1E-H, Fig. S3). This analysis also revealed a distinction in expression patterns of MX specific reporter lines with some genes showing expression exclusively in the outer or inner MX cell file (Fig. 1F/G, Fig. S3). Therefore, MX-specific cells were divided into clusters containing outer MX cells (cluster 3, 8, 12) or inner MX cells (0, 2, 4, 6) (Fig. 1C). Thus, despite their morphological similarities, our analysis indicates that outer and inner MX cell files display a divergent transcriptomic signature; suggesting these could be considered as different sub cell types. All scRNA-seq data is publicly available and integrated in our interactive browser (www.single-cell.be/plant).

Next, expression levels were calculated for each gene over each of the three different xylem sub cell type trajectories (PX, IMX and OMX) by means of the tradeSeq algorithm [18] (Fig. 1D). This results in expression dynamics that can be compared between genes for each of the different lineages (see Materials and Methods for details) (Table S2). Plotting known regulators shows that the dataset covers the entire developmental trajectory of PX while marker genes of very late events of xylem development seem missing in MX (Fig. S2). This observation can be attributed in part to limitations in generating protoplasts from cells located in the center of the root, and spatiotemporal difference in primary xylem differentiation in the root meristem.

In summary, our xylem-specific scRNA-seq dataset reveals distinct transcriptional identities for each of the xylem subtypes in the Arabidopsis primary root meristem and allows to follow pseudotime expression of genes across the developmental trajectories of all subtypes with great resolution.

### GRN analysis identifies ZHD3 and ZHD4 as regulators of PX development

In a pursuit to extract information about regulatory networks controlling xylem development from the xylem-specific scRNA-seq dataset, we applied MINI-EX (Motif-Informed Network Inference based on single-cell EXpression data) version 3.2 [19,20]. This gene regulatory network (GRN) analysis infers cell-type-specific networks and prioritizes the most functionally relevant regulators to unravel the complex interplay of transcription factors (TFs) controlling biological and developmental processes [19,20]. To assess the reliability of this unbiased analysis, we first evaluated how well known regulators are recovered. Indeed, many well-established regulators are recovered spanning from early establishment of the vascular pattern (*TMO3* and *TMO5* [21,22]), over initiators of differentiation like *VNDC, VND7* [4–7] and their upstream regulator *GATA12* [23] to ZAT-family transcription factors which control PCD onset downstream of *VND7* [24] (Fig. 2A-C). Besides these, positive regulators of xylem differentiation inhibitors such as *VNI1* are retrieved as well [25]. A GO-term enrichment of the top 50 candidates predicted by the GRN analysis reveals that processes associated with “regulation of secondary cell wall biogenesis”, “xylem development”, “phloem and xylem histogenesis” and “tracheary element differentiation” are amongst the most represented (Fig. 2C). This analysis confirmed that we captured central players of the entire PX developmental trajectory from initials to SCW deposition and PCD. For the MX trajectories, we did not capture the final steps of differentiation due to the protoplast isolation strategy used. As such, we decided to focus on the PX trajectory for further analysis.

**Figure 2.**
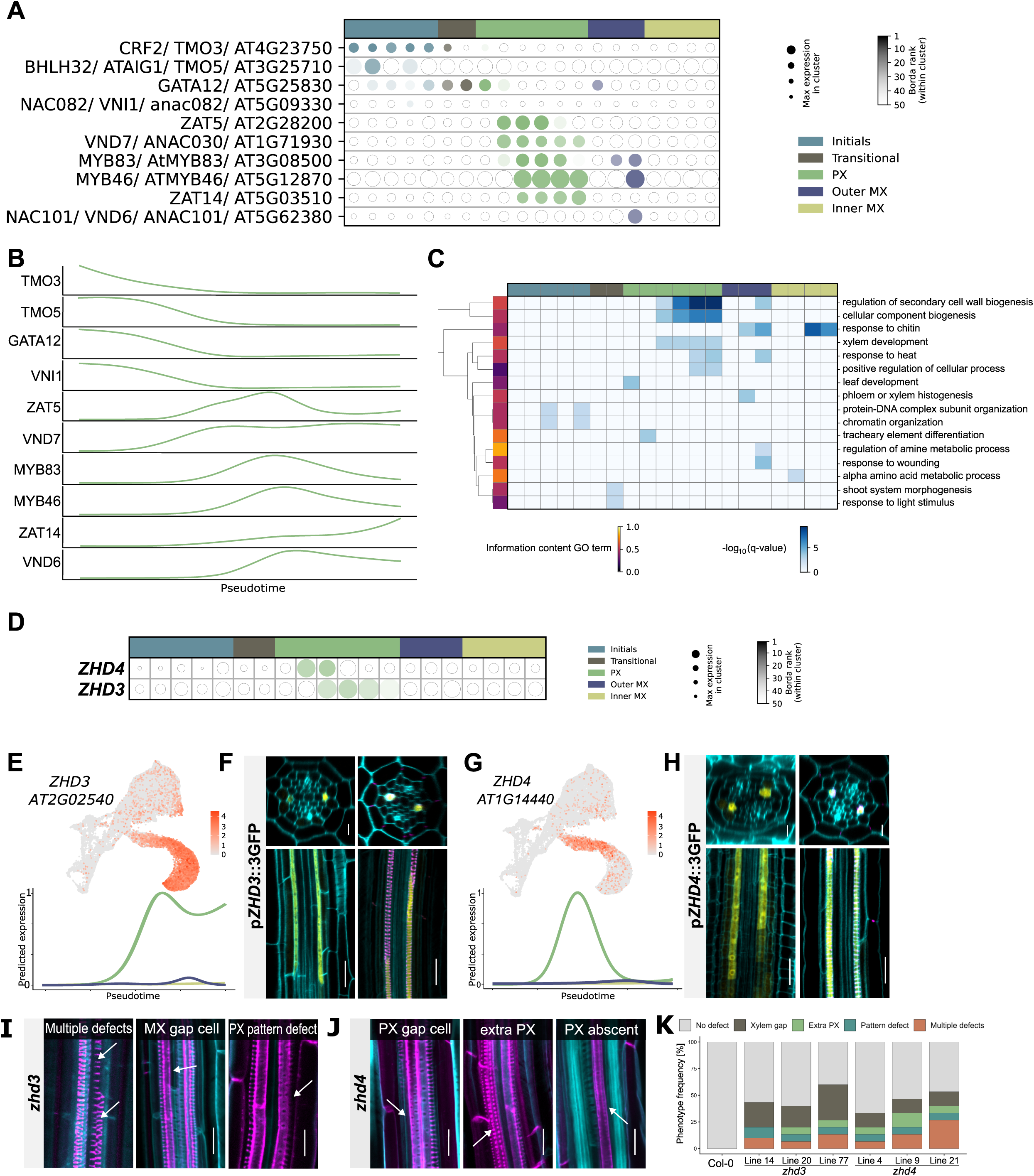
ZHD3 and ZHD4 are regulators of protoxylem development. **A.** Visualisation of the MINI-EX v3.2 output for known xylem regulators for each of the xylem cell lineages. **B.** Expression for these known regulators across pseudotime in our scRNA-seq dataset. **C.** GO term analysis output of the top 50 highest ranked TFs in the GRN analysis. **D.** Visualisation of the MINI-EX v3.2 output for the selected *ZHD3* and *ZHD4* regulators. **E-H**. Feature plot (top left), expression over pseudotime (bottom left) and confocal validation (right) showing both cross sections and longitudinal sections of the root apical meristem. Two positions in the root meristem are shown covering the longitudinal developmental zones. First appearance of the fluorescent signal is shown in the left image while differentiated PX is shown on the right. **I-J**. Confocal longitudinal images of the xylem defects observed in the *zhd3* and *zhd4* single mutants. **K.** Ǫuantification of the data represented in panels I-J. In panels F, H, I-J cell walls are counterstained using Calcofluo white (cyan), lignin is stained using basic fuchsin (magenta), and gene expression (p*GENE*::nGFP) in yellow. Scale bars are 20 μm.

Having established that this approach successfully identifies known regulators of xylem development, we next aimed to select novel putative regulators from its predictions. Amongst the 50 top-ranked TFs are two members of the *ZINC FINGER HOMEODOMAIN 3* (*ZHD3, AT2G02540*) and *ZHD4* (*AT1g14440*) that caught our attention due to their expression predicted to be specifically restricted to protoxylem cells (Fig. 2D-H) and their thus far unexplored role during primary vascular development. In order to validate the predicted expression pattern, we generated transcriptional reporter lines (p*ZHD3*::3GFP and p*ZHD4*::3GFP) and confirmed the protoxylem-exclusive expression for both genes (Fig. 2F, H). Relative expression levels along pseudotime show an earlier appearance of *ZHD4* transcripts compared to *ZHD3* during PX maturation (Fig. 2E, G), which was also confirmed in the reporter lines (Fig. 2 F, H). Within the *ZHD* family, other members have been recently reported to be involved in repressing cambium activity during secondary vascular growth [26]. However, *ZHD3* and *ZHD4* were predicted not to be expressed during secondary growth [27]. We confirmed this by analyzing expression of the reporter lines during this developmental stage (Fig. S4C, D). These analyses allowed us to rule out a putative role of *ZHD3* and *ZHD4* during secondary vascular development. In order to investigate the requirement of ZHD3 and ZHD4 for normal xylem development, we generated CRISPR-Cas9-mediated loss-of-function lines (Fig. S6 A-D). Although root length and architecture remained unaffected (Fig. S5A, B), a reduction of *ZHD3* or *ZHD4* activity induced a variety of developmental defects in mature xylem cells, including discontinuous xylem strands labeled as xylem gaps (Fig. 2I-K). Taken together, the GRN analysis prediction and validation by mutant analysis suggests *ZHD3* and *ZHD4* might act as regulators of protoxylem development in the primary root meristem.

### ZHD3/4 expression is sufficient to induce PX-like SCW depositions in any xylem cell file

To examine the function of *ZHD3* and *ZHD4* during xylem development in more detail, we expressed both TFs under the xylem-specific promoter of *AT3G42800* (labeled as *p*XYLEM) previously selected for validation of xylem initials (Fig. 1E) thus expanding the expression zone from PX to all five xylem files (Fig. 3 A,B, Fig. S5). This xylem-specific overexpression of both ZHD factors results in the appearance of helical SCW deposition in MX cell files (Fig. 3A,B, E) and an earlier appearance of SCW deposition, despite root length being unaffected (Fig. S6). Intrigued by this finding, we next investigated whether expression of *ZHD3* or *ZHD4* outside of the xylem domain would cause a global trans differentiation into PX-like cells; as has been observed for *VND* factors [28]. Driving *ZHD3* or *ZHD4* from the Ubiquitin promoter (p*Ubi*::XVE>>ZHD3/4) triggered the same PX-like SCW depositions in the MX cell files, but cells other than xylem remained unaffected (Fig. 3C, E; Fig. S5). Under endogenous conditions neither *ZHD3* nor *ZHD4* is expressed during secondary growth (Fig. S4C, D). However, since homologous genes are involved in secondary vascular development, we analyzed SCW depositions of secondary xylem vessels in induced p*Ubi*::XVE>>ZHD3/4. Contrary to what we observed during primary xylem formation, secondary xylem vessels retained a pitted SCW pattern (Fig. S4A, B). To further confirm that *ZHD3* and *ZHD4* are only capable to induce PX formation within the xylem axis, we expressed these TFs under a pericycle-specific promotor. Upon induction, no phenotype was observed (Fig. 3D, E, H; Fig. S5), supporting our hypothesis, that only cells primed to become xylem are capable to (trans-)differentiate into PX-like cells.

**Figure 3.**
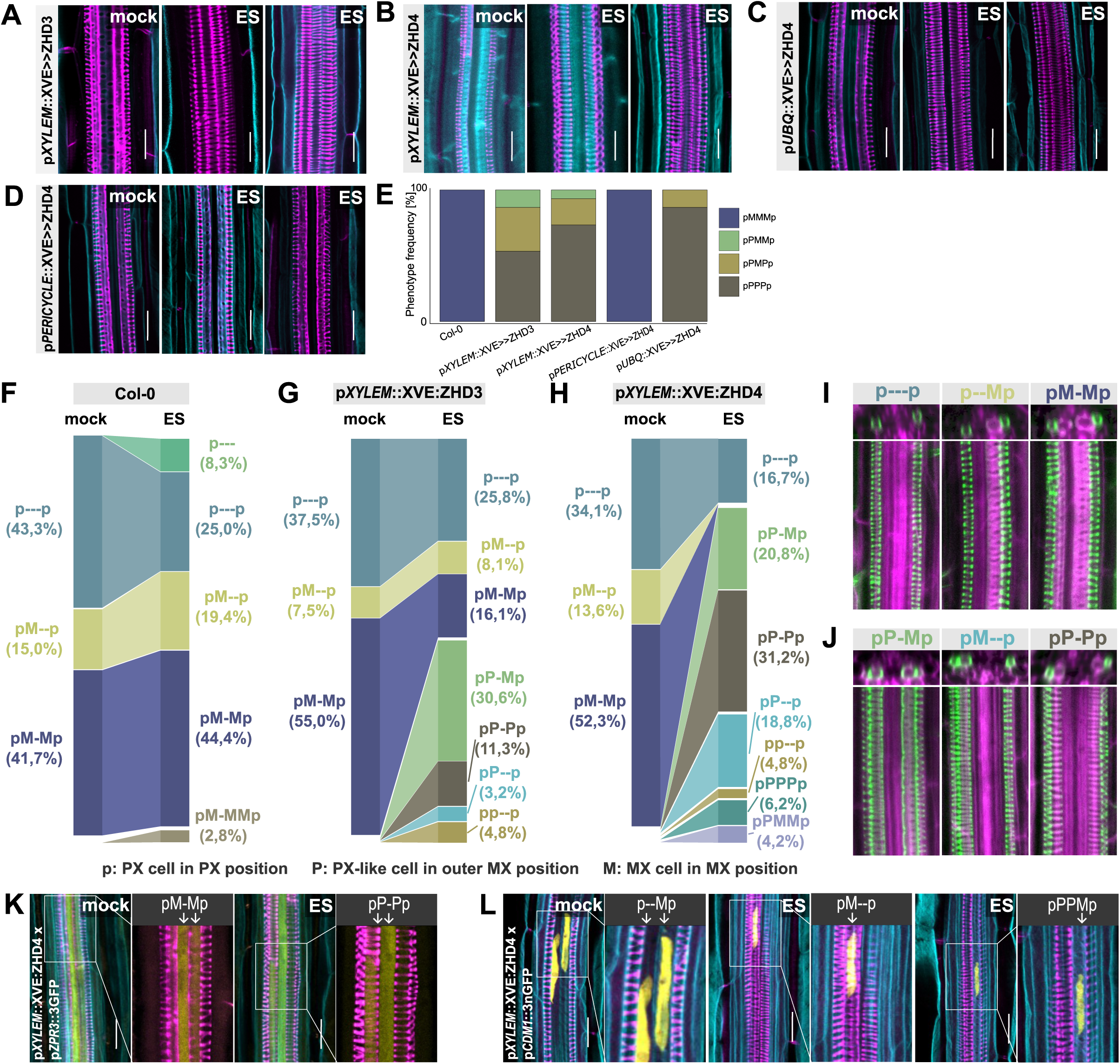
ZHD3/4 are sufficient to induce PX-like SCW depositions in xylem cells. **A-D.** Confocal longitudinal images of the xylem defects observed in the genotypes mentioned. **E.** Ǫuantification of the data represented in panels I-J. **F-H**. Detailed quantifications of xylem secondary cell wall morphology parameters in the indicated genotypes in mock and 10 μM estradiol (ES) conditions. **I-J**. Examples of the xylem phenotypes quantified in panels F-H. Panel I shows xylem development observed in Col-0 and panel J shows defects found in induced roots of p*XYLEM*::XVE>>ZHD3/4. **K-L**. Confocal longitudinal images of the xylem defects observed in the genotypes mentioned in mock and 10 μM estradiol (ES) conditions. In panels A-D and I-L, cell walls are counterstained using Calcofluo white (cyan), lignin is stained using basic fuchsin (magenta), and gene expression (p*GENE*::nGFP) in yellow. Scale bars are 20 μm.

The apparent formation of PX-like SCW depositions in the MX position could either be caused by a shift in identity from MX to PX; or by deposition of PX-like reinforcement of the cell wall in cells that maintain a MX identity. In order to discriminate between these scenario’s, we started by performing a more detailed examination of the observed phenotypes. Half of all induced p*XYLEM*::XVE>>ZHD3 roots and 83,3% of p*XYLEM*::XVE>>ZHD4 developed at least one strand with PX-like SCW depositions in the outer MX position, a defect that is never observed in Col-0 (Fig. 3F-H). Besides differences in SCW pattern, PX cells exhibit a smaller diameter compared to MX in control conditions. PX-like cells induced in the OMX position have an intermediate diameter while pitted outer MX and PX (in the PX position) cells maintain a size similar to control conditions (Fig. S6C). Furthermore, the width between individual cell wall bands of ectopic PX cells in *ZHD* mis-expression lines was reduced compared to PX cells in wild type (Fig. S6D). Moreover, PX-like cells in the MX position differentiated earlier than true outer MX cells in control conditions (Fig. S6E) as measured by comparing the distance from the root tip where the first xylem cell shows SCW cellulose deposition. Taken together, these detailed quantification of xylem cell parameters shows that the PX-like cells induced in the MX position show a clear divergence from the typical PX features. To further decipher the identity of ectopic PX cells, we thus investigated the expression of MX specific reporter genes from our single cell dataset in these PX-like induced cells (Fig. 4K,L). Concomitant with the intermediate morphological characteristics of the ectopic PX-like cells, those cells expressed one but not the other MX specific marker gene (Fig. 4K,L), suggesting there is no complete identity change occurring, but rather the activation of a specific part of the differentiation pathway. In summary, ectopic expression of *ZHD3* and *ZHD4* in all xylem cells results in a PX-like appearance of MX cells with features diverging from typical PX or MX features.

**Figure 4.**
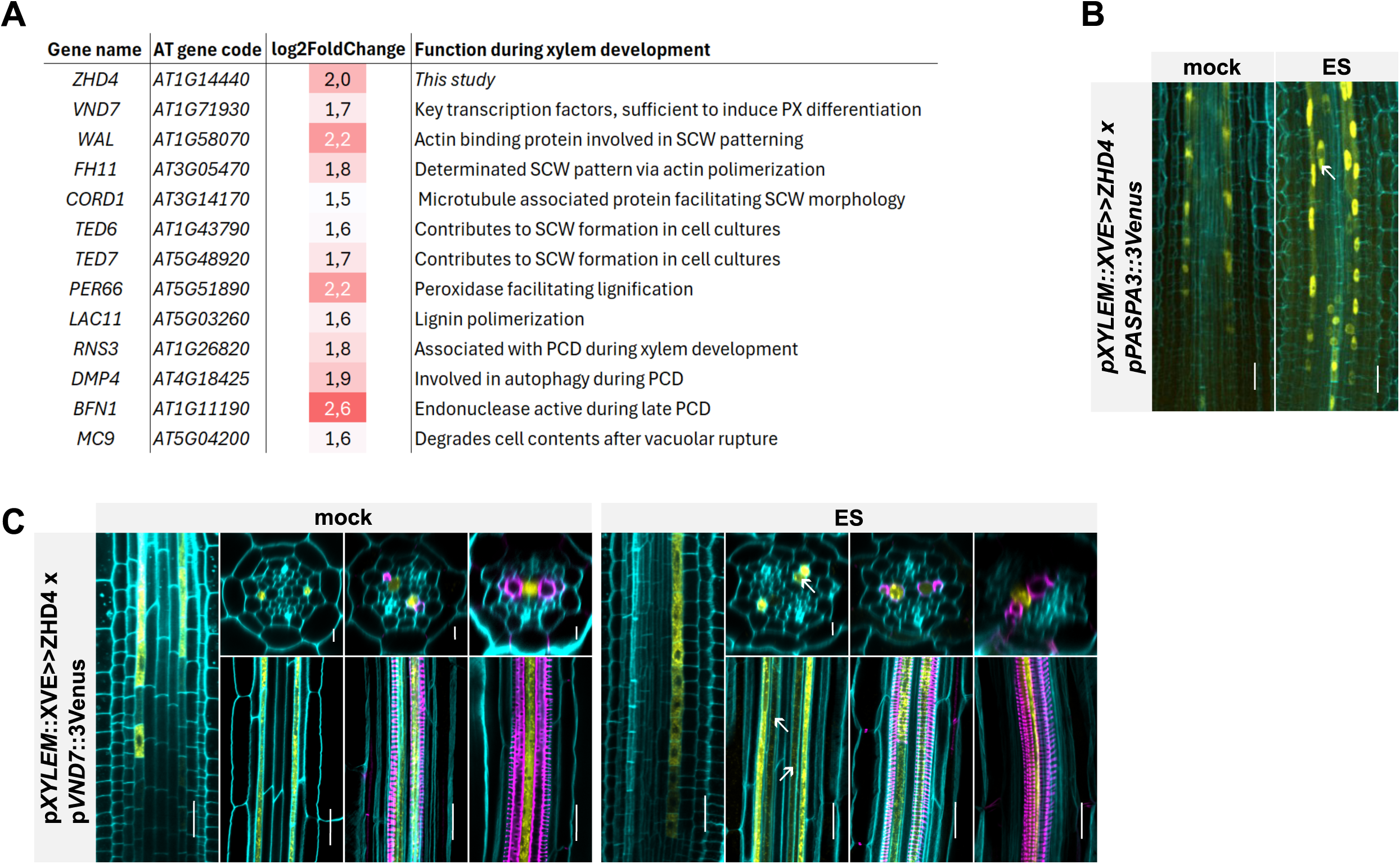
ZHD4 specifically activates the xylem differentiation machinery. **A.** Summary of the bulk RNA-seq data of ZHD4 induction showing significant differentially expressed genes. **B-D.** Confocal longitudinal images of the xylem defects observed in the genotypes mentioned in mock and 10 μM estradiol (ES) conditions. In C-D, both cross sections and longitudinal sections of the root apical meristem are shown with three positions in the root meristem covering the longitudinal developmental zones. In panels B-D cell walls are counterstained using Calcofluo white (cyan), lignin is stained using basic fuchsin (magenta), and gene expression (p*GENE*::nGFP) in yellow. Scale bars are 20 μm.

### ZHD4 specifically activates the xylem differentiation machinery

In order to understand the transdifferentiation phenotype observed upon expressing *ZHD3* and *ZHD4* in the MX domain at a molecular level, we next explored the transcriptomic changes that underlie the formation of PX-like cells in the outer MX position by performing a time series bulk RNA-seq experiment of *ZHD4* induction (Table S3). Many known regulators and executors of the xylem developmental machinery are significantly induced (Fig. 4A). The upregulated genes span a broad range of xylem developmental events from early specification of PX (*VND7* [28]) over actin and microtubule rearrangements (*WAL* [29], *FH11* [30]*, CORD1* [31]) and SCW deposition (*TEDC* and *TED7* [32]*, PERCC* [33]*, LAC11*[34] B) to execution of dPCD (*RNS3* [12]*, DMP4* [35]*, BFN1* [36]*, MCS* [37]) (Fig. 5A). Given *ZHD4* induces genes covering the entire developmental trajectory, this strongly points towards a role for *ZHD4* in transcriptionally activating the early steps of xylem development. Although no conclusions about the concrete mode of function can be drawn at this moment, *ZHD3* and *ZHD4* appear to be positive regulators of PX development with the ability to dictate a PX-like cell fate when expressed across the entire xylem axis.

To consolidate and visualize the outcome of the RNA-seq we introduced a marker for xylem specification (p*VND7*::3Venus), and PCD execution (p*PASPA3*::3Venus) in *p*XYLEM::XVE>>ZHD4 (Fig. 5B-D). This approach also allows us to distinguish between all three xylem subtypes and track changes in expression in PX, outer and inner MX. While *VND7* seems unaffected in PX, cells in the outer MX position show an earlier onset of expression (Fig. 4C). Expression of the PCD marker gene *PAPSA3* is intensified in PX and can be detected in cells located in the outer MX in the meristematic zone (Fig. 4 B). Together, our data suggest a role for *ZHD4* during the early events of xylem development; likely upstream of the known *VND7* master regulator.

### SCW depositions might be inadequate to determine cellular identity

Our functional characterization shows that misexpression of *ZHD3* or *ZHD4* is sufficient to induce a helical SCW pattern in all cell files of the xylem axis, but not in cells outside of the xylem axis. This suggests that other factors are required in order for the full differentiation process to occur or that a xylem cell identity (delivered through the ontogeny) is needed. Based purely on SCW morphology, those ZHD3/4 induced ectopic PX cells would be indeed labeled as PX, suggesting a switch in cellular identity. However, a more in-depth analysis of the PX-like cells in MX position shows that these cells 1) show measurable differences in SCW pattern compared to true PX cells and 2) express MX-specific marker genes. Thus, the ontogeny of the outer MX trajectory can therefore not be fully overwritten by a PX-like, helical SCW deposition. Similarly, despite the indistinguishable, pitted SCW pattern of outer and inner MX, we found that both cell types have divergent transcriptomic landscapes. Thus, assigning a cell identity based on morphological appearance of the xylem cell, as it has historically been practiced, might not be sufficient to accurately describe the cellular state of these xylem cells. Our data indicates that, at least for ZHD3/4 induced PX-like morphology, the cells likely retain their original MX identity. As such, it seems like a more detailed physiological analysis, ideally paired with a transcriptional approach is needed to confidently assign tissue identities.

## Materials and Methods

### Plant growth conditions

*Arabidopsis thaliana* ecotype Columbia 0 (Col-0) was used for all transformation procedures and phenotype analyses. After sterilization, seeds were stratified for 3 days at 4 °C and grown on half strength Murashige and Skoog (½ MS; Duchefa) medium [38] with or without 10μm Estradiol (ES). Seedlings and mature plants were cultivated at 22 °C under continuous light conditions. Plates were positioned vertically in the growth chamber. For measurements of xylem morphology parameters, seedlings were grown for 5 days on ½ MS medium without sucrose in absence (-ES) or presence (+ES) of 10 µM estradiol at 18 °C night and 23 °C day temperature in a 16-hour long-day light regime providing 110 µE. For averaging out light and temperature gradients, the seedlings were grown on a smart plate imaging robot while their growth trajectory was monitored [39].

### Protoplasting conditions and FACS

Three reporter lines with overlapping expression in the root primary xylem pTMO5::nTdT, pACL5::nGFP, and pATHB8::nYFP were used for fluorescence-activated cell sorting (FACS) in this study. Each reporter line was sorted and processed independently. Protoplasting was performed as previously described [17,40] with minor modifications. Briefly, root tips from 6-day-old seedlings were excised and incubated for approximately 1h at room temperature in protoplasting Solution B, comprising 1.5% (wt/vol) cellulysin and 0.1% (wt/vol) pectolyase dissolved in Solution A (600 mM mannitol, 2 mM MgCl₂, 0.1% [wt/vol] BSA, 2 mM CaCl₂, 2 mM MES, and 10 mM KCl, pH 5.5). Digestion conditions were optimized to preferentially dissociate inner vascular cells, even at the expense of outer cell layers, as assessed by periodic microscopic inspection. Following digestion, cells were filtered through a 70 μm cell strainer and centrifuged at 200 × g for 6 min. The pellet was resuspended in Solution A, passed through a 40 μm cell strainer, and stained with 4′,6-diamidino-2-phenylindole (DAPI) at a final concentration of 14 μM for live/dead discrimination. Cell sorting was performed using a BD FACS Aria II, and DAPI-negative protoplasts were collected for downstream analysis.

### 10X Genomics sample preparation, library construction, and sequencing

Sorted cells were centrifuged at 400×g for 6 min at 4°C and resuspended in Solution A to a final concentration of approximately 1000 cells/μl. Single-cell suspensions were loaded onto a GemCode Single Cell 3′ Gel Bead and Library Kit (v2 chemistry, 10x Genomics) or a Chromium Next GEM Single Cell 3’ GEM, Library C Gel Bead Kit (v3.1 chemistry, 10x Genomics) following the manufacturer’s instructions. Sequencing libraries were generated and sequenced on an Illumina NovaSeq 6000 platform according to 10x Genomics guidelines at the VIB Nucleomics Core Facility (VIB, Leuven). Raw data processing, including generation of the gene expression matrix, as well as downstream data analyses (clustering, cell type identity assignment, differential gene expression analysis, and quality control), were performed in coordination with the Single Cell Core Facility at VIB and Ghent University. Data comparison and development of an online visualization tool were also carried out in collaboration with the same facility.

### ScRNA-seq Dataset Analysis

Raw fastq files from four individual samples were mapped in parallel to the Ensembl TAIR10.40 Arabidopsis thaliana reference genome using 10x Cell Ranger (version 3.1.0) software cellranger count. For each sample, the resulting gene-to-cell count matrix was filtered for unique molecular identifier count, gene count, percentage mitochondrial genes and percentage of chloroplast genes using the number of mean absolute deviations (NMADs; lower limit: 1 NMAD, upper limit: 4 NMADs). For subsequent processing steps, the Seurat suite (version 4) was used [41]. Gene counts were normalized using SCTransform [42], and samples were integrated using the Seurat integration algorithm [20] . For both steps only the top 3000 variable genes were considered. Next, cells were clustered using the Louvain algorithm with a resolution of 0.8 and the first 25 PCs and, following UMAP dimension reduction and marker-based cell-type annotation, a selection comprising 28,622 cells and 22 cell-type clusters was made, which concluded the initial processing steps of the data set. Additionally, two non-annotated cell clusters were deemed uninformative and removed from further analyses. With this final selection of cells, samples were reprocessed as described above, starting from the filtered gene-to-cell count matrices and without any restrictions on the number of genes included. Cell-type clusters were delineated using the more recent Leiden algorithm, with a resolution of 0.6 and the first 50 PCs, implemented in the python package SCANPY.

Cell-type-specific GRNs were inferred using MINI-EX version 3.2 with default parameters and using the standard (“std”) ranking procedure, resulting in regulator prioritization that is independent of gene ontology (GO) information. Raw RNA expression counts were used as the input matrix, and differentially expressed genes used for GRN inference were generated from log-normalized RNA counts utilizing Seurat’s (version 5.1.0) FindMarkers (Wilcoxon test, default parameters) in R (version 4.2.2). For each cluster, the resulting top 50 (based on the MINI-EX “ClusterRank”) cluster-specific regulators were tested for GO enrichment using a hypergeometric test with Benjamini-Hochberg correction (using the enricher tool of MINI-EX version 3.2 at https://github.com/VIB-PSB/MINI-EX/blob/main/bin/enricher_v.3.3.1). GO information was obtained from MINI-EX version 3.2 (https://github.com/VIB-PSB/MINI-EX/blob/main/data/ath/ath_full_BP_expcur_ext_names.tsv). The gene set used as the enrichment background consisted of all expressed transcription factors (TFs; genes in MINI-EX’s Arabidopsis TF list that are present in the raw RNA expression matrix). For visualization, only highly significant GO terms with an adjusted p-value < 0.005 are shown, and redundant GO terms were removed by retaining only the most significant GO term across related (ancestral or descendent) terms, per cluster.

### Plasmid construction and transformation

The constructs were generated on the basis of the Golden Gate system, as indicated in Table S4. To generate reporter lines used for validation if the scRNA-seq atlas or localization of *ZHD3* and *ZHD4* expression domains, promoter regions (up to ∼3 kb) were amplified, introduced into the Golden Gate entry module (pGGA000) and recombined with nuclear or cytosolic GFP into the destination vector pFASTRK-AG [43]. For ES inducible lines of *ZHD3* and *ZHD4* under *pUBI, pAT3G42800* (labeled as *pXYLEM*) and *pPERICYCLE*, promotors were fused to and XVE module and coding regions amplified from gDNA. All entry modules were recombined into the destination vector pFASTRK-AG [43]. To generate CRISPR-Cas9 knockout mutants, protospacers targeting the exon region of the gene were designed using the CRISPR P tool [44] and checked for off target specificity using https://design.synthego.com/#/validate and https://chopchop.cbu.uib.no/. Protospacer sequences are listed in (Table S5). Oligos containing BbsI overhangs were annealed and ligated into the corresponding pEN Chimera entry vectors using Golden Gate cloning. Entry vectors were assembled into the pFASTRK_AtCas9_AG expression vector using Golden Gate assembly. All constructs were transformed into Agrobacterium tumefaciens C58C1 and introduced into Arabidopsis thaliana by floral dipping [45]. A schematic overview of the genomic changes introduced in *zhd3* and *zhd4* loss-of-function lines is shown in Fig. S6C, D.

### Histochemical staining and confocal imaging

Histochemical staining was performed on 7-day-old seedlings using the modified protocol as described in [46]. Seedlings were fixed with 4% Paraformaldehyde (PFA), followed by 2 washes in phosphate-buffered saline (PBS). Tissue clearing was performed using the ClearSee solution overnight, followed by the lignin staining using 0.2% Basic Fuchsin dissolved in ClearSee overnight, and cellulose staining using 0.1% Calcofluor White dissolved in ClearSee for a maximum of 5 minutes. We imaged Basic Fuchsin using 561-nm excitation with detection at 600–650 nm, and Calcofluor White using 405-nm excitation with detection at 425–475 nm using the Leica Stellaris 5 confocal microscope.

### Ǫuantification of SCW characteristics and statistics

Seedlings were fixed in 4% paraformaldehyde, washed in PBS, cleared using the ClearSee method, and stained with Calcofluor White and Basic Fuchsin [46]. Roots were imaged on an LSM880 microscope equipped with an Airyscan detector and a 40× water-immersion objective (NA 1.2). Calcofluor White was excited at 405 nm and detected over 411-459 nm. Basic Fuchsin was excited at 561 nm and detected over 600-650 nm (main beam splitters MBS −405 and MBS 488/561; confocal pinhole 0.93 and 0.65 Airy units, respectively). Voxel size was 208 × 208 × 300 nm (8-bit). The lateral resolution limit is therefore approx. 0.22 µm in the Calcofluor channel and 0.32 µm in the Fuchsin channel; measured band widths (approx. 0.9 µm) are approximately three times the point-spread function, so absolute band and gap widths include a point-spread broadening component. This component is constant across groups and cannot create or mask a between-group difference, but absolute values should be read as upper bounds. Xylem secondary wall patterns were analysed with RadialJ, an unpublished Fiji/ImageJ plugin Confocal z-stacks were resampled to isotropic voxels (200 nm) and reoriented so that the root’s growth axis was perpendicular to the image plane, which gives cross-sectional views of the xylem. Protoxylem and metaxylem vessels were identified by position and wall signal, and the approximate centre of each vessel was marked with a single mouse click. From these seeds, vessel outlines were segmented in the first slice by marker-controlled watershed, using a weighted combination of the cellulose and lignin channels. Segmentation was propagated slice by slice along the growth axis, yielding the outline and centroid of each vessel over its full length. Each vessel was then reprojected into a vessel-centred reference frame: wall signal was sampled radially (0-360°) around the centroid at every axial position, giving (i) a radial projection of angle φ against axial position z and (ii) an unrolled projection, in which signal is plotted against true arc length s along the vessel circumference. The unrolled projection preserves the true surface geometry and therefore reflects differences in vessel diameter.

Wall pattern parameters were measured on these projections. Diameter and circularity were taken from the segmented outline at every axial position in which the vessel was resolved and averaged. Bands were detected by peak finding along 100 random 25 µm line profiles per vessel oriented along the growth axis; gaps were detected by applying the same procedure to the inverted signal. This yielded band width and gap width. Per-vessel means of these measurements are the values plotted. Vessels were classified by their position in the five-cell xylem axis, the two protoxylem poles, the two outer metaxylem positions and the central metaxylem position, together with their wall pattern type. Protoxylem (p) vessels showed banded wall patterns at a protoxylem pole; intermediate (P) vessels showed banded wall patterns at a metaxylem position; metaxylem (M) vessels showed pitted wall patterns at a metaxylem position. Vessels for which neither position nor pattern could be assigned unambiguously (type U, 17 of 634) were excluded from all analyses. Band and gap parameters are defined only for banded wall patterns and were therefore measured on p and P vessels only (Fig. S6).

For statistical analysis of this data: because several vessels were measured per recording and several recordings per seedling, vessel-level values are not independent and were analysed with linear mixed models rather than treating vessels as replicates. For each figure a model of the form measurement ∼ group + (1|seedling) + (1|recording) was fitted by restricted maximum likelihood (REML) to the vessel-level data of the groups shown in that panel; where the recording-level variance was estimated at zero (Fig. S6) the model was reduced to a seedling random intercept. Denominator degrees of freedom were obtained by the Kenward–Roger approximation, and groups were compared by Tukey-adjusted pairwise contrasts of estimated marginal means. Letters in the panels were derived from those adjusted p-values. Analyses used R 4.5.3 with lme4 2.0.6, lmerTest 3.2.1, pbkrtest 0.5.5 and emmeans 2.0.4; letter displays used multcompView 0.1.12.

Comparisons between vessel types within a genotype are made within the same seedlings and are therefore paired, carrying approx. 200-320 denominator degrees of freedom. Comparisons between genotypes or between ±ES rest on 3-5 seedlings and carry approx. 9-20. Statistical resolution is consequently set by the number of seedlings, not the number of vessels. With the present design the Tukey-adjusted smallest detectable difference is approximately 1.1–1.5 µm for metaxylem diameter (Fig. S6), 0.36 µm for protoxylem diameter (Fig. S6), 0.08 µm within and 0.25 µm between genotypes for gap width (Fig. S6), and 0.04 µm within and 0.11 µm between genotypes for band width (Fig. S6). Non-significant results in Fig. S6 should therefore be read as excluding effects larger than these values, not as excluding any effect.Robustness checks: vessel-level variance differed about 3.4-fold between the groups of Panel 3 (Levene-type test p = 8 × 10-23). Refitting that panel with group-specific residual variances (nlme 3.1.170, varIdent) left 26 of 28 pairwise verdicts unchanged. The two that changed both involve ZHD4 +ES M, which rests on 4 vessels from 2 seedlings and should be treated as indicative only. Model assumptions were not otherwise assessed.

For vessel phenotype mapping along the root (Fig. S6), each root was imaged at successive positions along its length, and the cumulative distance between consecutive fields of view was recorded (median step 335 µm; 290 positions from 24 roots). Recording began at the first position at which a patterned secondary cell wall was detected on the camera, which defines 0 µm; distance increases shootward and is not referenced to the quiescent centre or the columella. At each position the differentiated vessels across the stele were scored as a five-character map (the two protoxylem poles, the two metaxylem files and the central file), with an unassigned character where no secondary cell wall had formed. Positions were grouped into 500 µm intervals; this width was chosen because, at the observed median step, 90% of intervals within a root’s imaged range contain at least one field of view, against 66% for 250 µm intervals. For each interval and vessel type, incidence was calculated as the number of roots in which at least one vessel of that type was observed divided by the number of roots imaged in that interval, so the three types are independent incidences and do not sum to one. Type presence was not carried forward between positions: appearance along the root is close to monotone but not strictly so (6 reversals in 290 transitions, mostly of the intermediate class), and imputation would overstate the intermediate class. With four roots per genotype and treatment, incidence takes only the values 0, 0.25, 0.50, 0.75 and 1.00; interval-by-interval counts are tabulated in Table S6.

### RNA-seq data processing and bioinformatic analysis

5-day-old plants of pXYLEM::XVE>>ZHD4 were transferred to ½ MS plates containing 10 μM Estradiol or to DMSO mock plates and sampled at 8h and 24h. Approximately 300 individual root tips were collected per sample replicate, and three biological repeats per time point were used. RNA extraction was performed using the TRIzol-based RNA isolation protocol. Ǫuality control for RNA samples was performed using the Agilent TapeStation system. For further processing samples were sent to BGI for library preparation and RNA sequencing. Raw FASTǪ files with paired-end RNA-seq reads were subjected to quality assessment using FastǪC (v0.11.9), and quality reports were summarized with MultiǪC (v1.33). Low-quality bases and short reads were removed using Trimmomatic (v0.39) with the following parameters: leading:3, trailing:3, slidingwindow:4:5 and minlen:80. Filtered reads were aligned to the Arabidopsis thaliana TAIR10 reference genome using HISAT2 (v2.2.1). In addition, the first 10 nucleotides from the 5′ end of each read were removed during alignment using the –trim 5 10 option. Resulting alignments were sorted and converted to BAM format using SAMtools (v1.13). Gene read counts were generated with featureCounts (v2.0.3) using the TAIR10 gene annotation (release 62).

Differential expression analysis was carried out using DESeq2 version 1.52.0. Log2FC values were adjusted to account for low-count genes using lfcShrink, and genes with absolute adjusted Log2FC values > 1.3 and adjusted p-value < 0.05 were selected as differentially expressed genes.

## Data availability

The scRNA-seq data described in this study are available in NCBI GEO under accession GEO: (will be added upon submission). These data will be publicly available and integrated in our interactive browser upon acceptance of the manuscript (https://single-cell.be/plant). The bulk RNA-seq data described in this study are available as NCBI GEO (will be added upon submission).

## Acknowledgements

We are thankful to the VIB Single Cell Core, VIB Flow Core Ghent and VIB Nucleomics for support and access to the instrument park (vib.be/technologies). T.E was supported by the European Research Council (ERC) CoG PIPELINES grant (101043257); M.S-S. was funded by the Ghent University Special Research Fund (BOF20/GOA/012). C.V.D.M. received financial support by the Deutsche Forschungsgemeinschaft (DFG, German Research Foundation – project number: 527041612) and the Research Foundation – Flanders (FWO grants 1221424N, G000822N and G0D0515N). R.S. was funded by Deutsche Forschungsgemeinschaft (DFG) grants (453188536, 512328399). A.S. was supported through association with the International Max Planck Research School for Molecular Plant Science (IMPRS MolPlant).

## Author contributions

Conceptualization: CvdM and BDR; Data Curation: AGN, CvdM, JS, TD, TE, MSS, RS; formal analysis: all authors; Funding Acquisition: CvdM, RS, KV and BDR; Investigation: all authors; Supervision: CvdM, RS, KV and BDR; Visualization: CvdM, JS, TD, TE, RS; Writing: CvdM and BDR, Reviewing C Editing: All authors.

